# Enzyme Kinetics and Inhibition Parameters of Human Leukocyte Glucosylceramidase

**DOI:** 10.1101/2020.02.23.961599

**Authors:** Mesut Karataş, Şenol Doğan, Emrulla Spahiu, Adna Ašić, Larisa Bešić, Yusuf Turan

## Abstract

Glucosylceramidase (GCase) is a lysosomal enzyme that catalyzes the cleavage of β-glucosidic linkage of glucocerebroside (GC) into glucose and ceramide; thereby, plays an essential function in the degradation of complex lipids and the turnover of cellular membranes.

The growing list of 460 mutations in the gene coding for it—glucosylceramidase beta acid 1 (*GBA1*)—is reported to abolish its catalytic activity and decrease its enzyme stability, associating it with severe health conditions such as Gaucher disease (GD), Parkinson Disease and Dementia with Lewy bodies.

Although the three-dimensional structure of wild type glucosylceramidase is elucidated, little is known about its features in human cells. Moreover, alternative sources of GCase that prove to be effective in the treatment of diseases with enzyme treatment therapies, impose the need for simple and cost-effective procedures to study the enzyme behaviour. This work, for the first time, shows a well established, yet simple, cost- and time-efficient protocol for the study of GCase enzyme in human leukocytes by the artificial substrate PNPG. Characterization of the enzyme in human leukocytes for activation parameters (optimal pH, Km, and Vmax) and enzyme inhibition, was done. The results indicate that the optimum pH of GCase enzyme with PNPG is 5.1. The K_m_ and V_max_ values were 12.6mM and 333 U/mg, respectively. Gluconolactone successfully inhibits GCase in a competitive manner, with a K_i_ value of 0.023 mM and IC _50_ of 0.047 mM. Glucose inhibition was uncompetitive with a K_i_ of 1.94 mM and IC_50_ of 55.3 mM. This is the first report for the inhibitory effect of glucose, δ-gluconolactone on leukocyte GCase activity.

## Introduction

Glucosylceramidase (GCase, EC 3.2.1.45) is a lysosomal enzyme with 497 amino acids and three discontinuous domains [1]. It is a β-glucosidase enzyme that catalyzes the cleavage of glucosylceramide (GC) into glucose and ceramide [2]; thereby, it plays a vital function in the degradation of complex lipids and the turnover of cellular membranes [3]. The main cellular compartment of GCase is the lysosomal membrane [4]. The activator proteins Saposin A and C, synergistically facilitate the interaction between the lipid substrate and enzyme, resulting in increased GCase enzymatic activity [5, 6]. The substrate (GC) is synthesized from the degradation of glycosphingolipids in the membranes of apoptotic white blood cells and senescent red blood cells [7].

Gaucher disease (GD, OMIM#230800) is the most protracted health condition related to defective GCase activity [6]. The recent studies, however, further associated GCase with severe health conditions such as Parkinson’s Disease (PD, OMIM#168600) [8], Dementia with Lewy bodies (DLB, OMIM# 127750) [9] and colorectal cancer [10]. GCase is encoded by the housekeeping gene glucosylceramidase beta acid 1 (*GBA1*, OMIM*606463) with cytogenetic location 1q21 [11]. More than 460 mutations are reported to disrupt the *GBA1* gene [12, 13], thereby abolishing its catalytic activity [14] and decreasing enzyme stability [15]. The improper breakdown of highly hydrophobic GC results in its accumulation in the lysosomes of macrophages, leading to dysfunctions in spleen, liver, and bone marrow [16, 17]. The abnormal accumulation and storage of these substances damage tissues and organs, causing the clinical manifestations of GD [18], which is the most frequent lysosomal storage disorder [19, 20]. GD is shown to produce a continuum of different clinical conditions, from the fatal perinatal disorder to the asymptomatic one [21]. The efficiency of enzyme replacement therapy (ERT) in GD, has led to extensive studies carried out in alternative sources of GCase [22, 23]. GD has a global occurrence of 1:50,000 [24], but due to the founder effect, it shows a higher prevalence in Ashkenazi Jewish (Eastern European), Spanish, and Portuguese populations [25]. On the other hand, PD is a closely related neurological complication of GD, which is seen with carriers of heterozygous mutations in the *GBA1* gene being susceptible to developing it [8]. Recent genetic and pathological reports suggest *GBA1* mutations as critical risk factors for DLB as well [26].

Detection of GD by GCase assay is done in a variety of human tissues, including the liver [27], brain [5], and placenta [28]. Although any human tissue can be used to study GCase activity, in our study, we used human blood as the most readily available one. *In vitro*, GCase is solubilized from membranes by the extraction of lysosomal membranes using a detergent—TritonX-100 [29–31]. The non-ionic detergent Triton X-100 removes the endogenous natural lipid activators that are found in lysosomal membranes [32], consequently rendering Glucosylceramidase inactive [29]. Therefore, the reconstitution of GCase activity requires the inclusion of an anionic detergent—sodium taurocholate [39–47]—in the assay medium. The detergent sodium taurocholate is an essential component of GCase assay and can be used with artificial substrates to measure the GCase activity with minimal interference from other β-glucosidases [30]. Sodium taurocholate inhibits nonspecific β-glucosidase, which cannot hydrolyze glucocerebroside and stimulate GCase activity [41]. Therefore a crude protein extract is suitable without the need for a purified protein. The lysosomal GCase, *in vivo*, is active on its natural lipid substrate glucosylceramide. *In vitro*, artificial substrates such as PNPG (para-Nitrophenyl-β-D-glucopyranoside) [46–48] and 4-MUG (4-methylumbelliferyl-β-D-glucopyranoside) [49–51] are very convenient ways of studying its activity due to a high yield of products that strongly absorb around 400 nm [53].

In this work, GCase enzyme activity in human leukocytes by using an artificial substrate is studied and for the first time the inhibitory effects of glucose and δ-gluconolactone are shown. A novel protocol was designed for brief biochemical characterization of activation parameters (optimal pH, Km, and Vmax) and enzyme inhibition.

## Materials and Methods

### Chemicals and samples

*p*-nitrophenyl-β-D-glucopyranoside (PNPG), pure sodium taurocholate (6 mg/ml), Triton X-100, glucose, and D-gluconolactone (D-glucono-1,5-lactone), were obtained from Sigma-Aldrich (Germany). Additionally, we used pure sodium acetate buffer (50 mM; 0.4355% w/v CH_3_COO Na/, 0.1089% w/v CH_3_COOH, pH 5), and lysis buffer (155 mmol/L NH_4_Cl; 10 mmol/L NaHCO_3_; 0.1 mmol/L EDTA) obtained from Semikem (Bosnia and Herzegovina). Blood samples were acquired using the guidelines and approval of the University of Burch Research Ethics Committee (30052016). Written informed consent was obtained from two healthy adult male donors. Whole EDTA blood (10 ml) was collected aseptically.

### Isolation of Leukocytes

Leukocytes were isolated by centrifugation after specific lysis of erythrocytes, according to the method of Peters et al. [42]. Aseptically collected whole EDTA blood (10 ml) was centrifuged at 1500*g* at 4°C for ten minutes, followed by the removal of plasma. The original blood volume was restored with 0.9% (w/v) NaCl solution and the blood suspension was transferred to a 50 ml conical centrifuge tube, followed by the addition of 40 ml cold hypotonic lysis buffer. The tube was left to stand on the ice and occasionally mixed for approximately ten minutes. The suspension was centrifuged again at 1500*g* at 4°C for five minutes. Leukocytes formed a small, firm pellet on the side of the tube while the erythrocyte ghosts formed a large loose pellet at the bottom of the tube. Both the hemoglobin-containing supernatant and the loose red cell ghosts were removed by aspiration using a Pasteur pipette. Leukocyte pellet was suspended in 5 ml cold hypotonic lysis buffer. The tube was left to stand on the ice. After 10 minutes, the cell suspension was diluted up to 50 ml with cold 0.9% (w/v) NaCl solution and centrifuged again at 1500*g* at 4°C for five minutes. The supernatant was discarded, and the leukocyte pellet was suspended in 10 ml 0.9% (w/v) NaCl solution followed by centrifugation at 1500*g* at 4°C for ten minutes. The supernatant was removed entirely, and samples were stored at −80°C for further use. A step by step protocol is uploaded at protocols.io (dx.doi.org/10.17504/protocols.io.bcv2iw8e) [54]

### Extraction of GCase from leukocyte pellet

GCase extraction protocol is modified from Kara et al. [55]. Leukocyte cell pellets equivalent to 10 ml of whole blood were thawed and suspended in 1.0 ml of 0.1% (v/v) detergent TritonX-100, and subjected to six cycles of freezing-thawing by alternating tubes, at approximately three-minute intervals, using a dry-ice/ethanol bath and water bath (−70°C /27°C). The total homogenate was centrifuged at 2100*g* for 5 minutes at 10°C and supernatant was put into a clean tube in an ice bath for assay. Protein concentration was estimated by the Bradford method, using bovine serum albumin as standard [56].

### Leukocyte Glucosylceramidase Assay

Glucosylceramidase activity was determined spectrophotometrically using 140 μl assay mixtures. The standard reaction mixture consisted of 70 μl of enzyme solution and 70 μl of 5 mM PNPG substrate in 50 mM sodium acetate buffer (pH 5.0), sodium taurocholate was dissolved in the substrate solution [30]. After incubation at 37°C for 45 min, the reaction was terminated by the addition of 70 μl of 0.5 M sodium carbonate (Na_2_CO_3_, pH 10.3). Absorbance was read at 400 nm against the blank solution containing substrate buffer instead of the enzyme extract. One unit of GCase activity was defined as the amount of enzyme required to release one μmol of p-nitrophenol from PNPG per minute (1 U), at 37°C and pH 5.0.

### In vitro Inhibition Studies and Determination of Kinetic Parameters

All the samples were assayed in the presence of pure sodium taurocholate at pH 5.0. The substrate concentrations for each sample were: 0.71, 1.07, 1.43, 1.79, and 2.50 mmol per liter. Incubations were carried out at 37°C for 45 min. The K_m_ and V_max_ were obtained through the Lineweaver-Burk plot, whereby the best fit lines were calculated by linear regression analysis. Inhibition experiments were performed using PNPG as substrate, and different final concentrations of δ-gluconolactone, glucose as possible inhibitors. A double reciprocal Lineweaver–Burk plot was used to calculate the parameters. The activity of the β-Glucosylceramidase at four different concentrations of each inhibitor was determined by regression analysis. Results are expressed as % activity, assuming 100% enzyme activity in the absence of an inhibitor. In the presence of possible inhibitors, % activity was calculated by using the following equation: % activity = 100−[(A_o_− A_i_)/A_o_)] ×100.

where A_o_ is the initial GCase activity (without inhibitor), and A_i_ is the GCase activity with the inhibitor. The inhibitor concentration that reduces the enzymatic activity by 50% (I_50_ values) was determined from the plots.

## Results and Discussion

### GCase Extraction and Optimization of Stability

Protein studies require proper preservation of proteins in order to work with the same quality of protein throughout the steps [57]. We performed the optimization study for the stability of GCase in leukocyte homogenate in the course of storage in −20°C frost-free freezer. It was observed that the activity of GCase, exhibits only minor changes after ten and twenty days of storage, in the presence of pure sodium taurocholate (Figure A1). In our study, GCase present in the crude protein extract was partially purified. However, the fact that cleavage of PNPG substrate is only possible by GCase, makes our way a convenient one to study the enzyme kinetics of GCase. Furthermore, UV-vis spectroscopy with an absorbance signal higher than 320nm is widely used to study protein preparations and monitoring enzyme activity [58]. In our case, the product of the catalyzed reaction, p-nitrophenol, was continuously released from PNPG, and this product strongly absorbed light at 400 nm [53]. With this theoretical framework, we developed a simple, cost- and time-efficient technique to study the activity of this enzyme with high importance in many diseases and industry.

### Determination of Optimal pH

The conformational stability and catalytic activity of enzymes are in direct relationship to their pH [59]. To determine the optimal pH value for the hydrolysis of PNPG by leukocyte GCase, we examined the effects of variable pH (4.0-5.5) at a constant substrate concentration (5mM). Figure 1.A shows the pH activity curve of leukocyte GCase assayed in the presence of sodium taurocholate. Leukocyte GCase was optimally active at pH 5.1, after which the enzyme activity started to decline (Figure 1.B). The optimal pH of GCase activity from different sources and different substrate is reported to be in 4.7–5.9 intervals [60], which is consistent with its lysosomal function and our finding.

**Figure 1.**
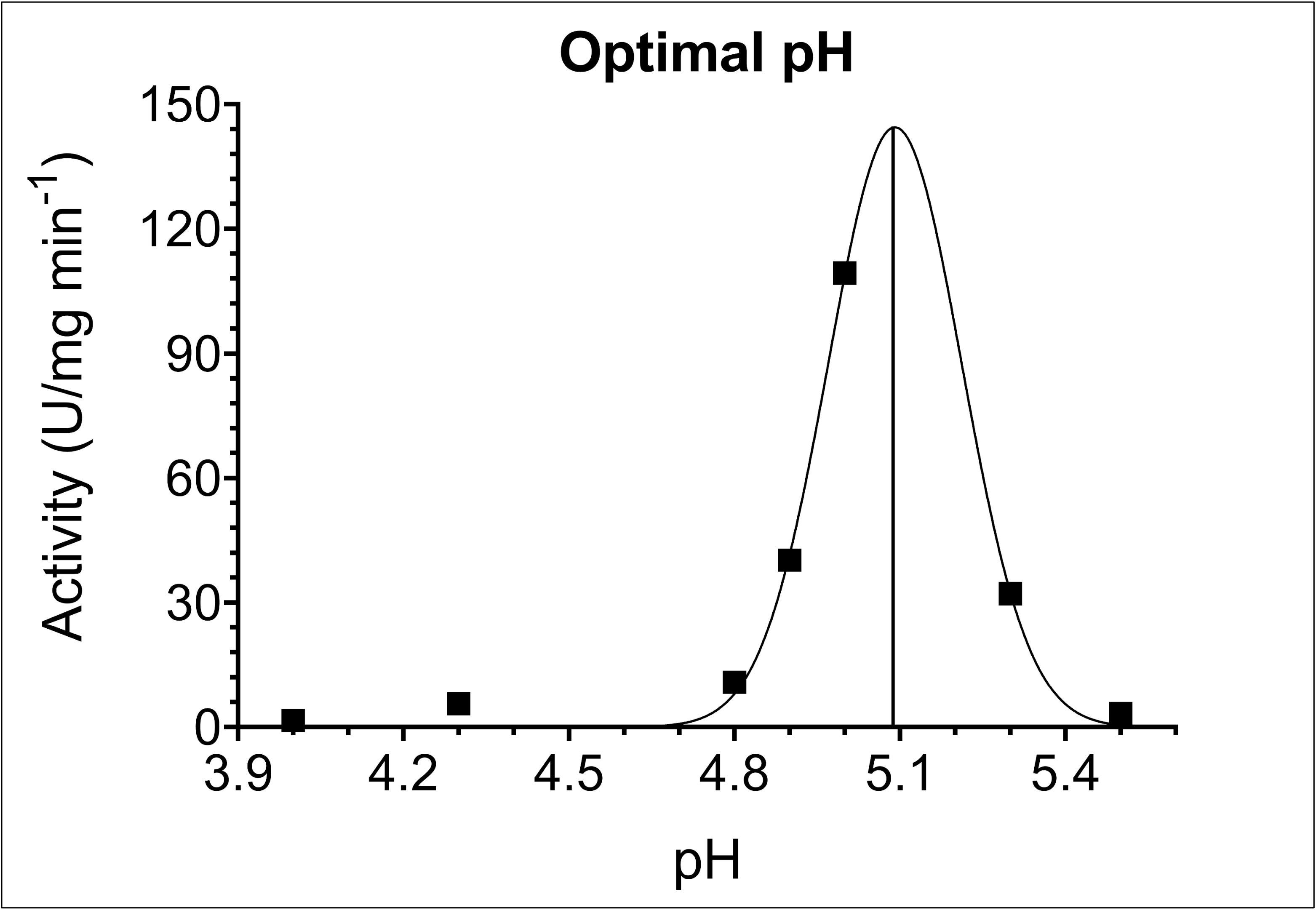
(**A**) The optimal pH for Human Leukocyte Glucosylceramidase (GCase) activity, is measured to be 5.1. The influence of varying pH values on the activity was determined by using sodium acetate buffer in the pH range of 4.0 to 5.5. (**B**) Lineweaver-Burk plot for Michaelis constant determination of GCase for different *p*-nitrophenyl-β-D-glucopyranoside (PNPG) substrate concentrations (0.71 mM-2.50 mM) in sodium acetate buffer (50mM, pH 5.0), reveals a K_m_ of 12.6 mM and V_max_ of 333 U/mg. All samples were assayed in the presence of 6 mg / ml sodium taurocholate and 70 μl of enzyme solution.

### Determination of Glucosylceramidase Km and Vmax for PNPG substrate

The Michaelis constant (K_m_) is a measure of the affinity of the enzyme towards the substrate, with smaller values representing higher affinity. The Michaelis constant (K_m_) and the maximum rate (V_max_) of leukocyte GCase were obtained through the Lineweaver-Burk plot (Figure 1.B) with artificial substrate PNPG in concentrations ranging from 0.71 mM to 2.50 mM. K_m_ and V_max_ values were found to be 12.6 mM and 333 U/mg, respectively, in reasonable agreement with previously reported (593 mM) studies [53].

### In vitro Enzyme Inhibitor Studies

The inhibition study was performed using the PNPG as a substrate. Inhibitors were defined to be gluconolactone and glucose. The results indicated that gluconolactone was the most effective inhibitor acting in competitively with a IC_50_ value of 0.047 mM (Figure 2.A) and K_i_ value of 0.023 mM (Figure 2B). The changing apparent K_m_ to higher values in higher gluconolactone concentrations is an indicator of competitive inhibition (Table 2). A report from placental cells has shown similar results [45]. However, as can be seen in Table 2, glucose was shown to be an uncompetitive inhibitor of PNPG hydrolysis, with a IC_50_ value of 55.3 mM (Figure 3A) and K_i_ value of 1.94 mM (Figure 3B). Based on literature, this is the first study to investigate gluconolactone and glucose inhibition in human leukocyte GCase. However, the inhibition kinetics of β-glucosidase (EC3.2.1.21) from several plants, fungi and especially microorganisms [62–66] has extensively been studied using glucose as an inhibitor, since glucose inhibition of β-glucosidase is undesirable if the enzymatic hydrolysis of cellulose is performed as an industrial process [67, 68].

**Figure 2.**
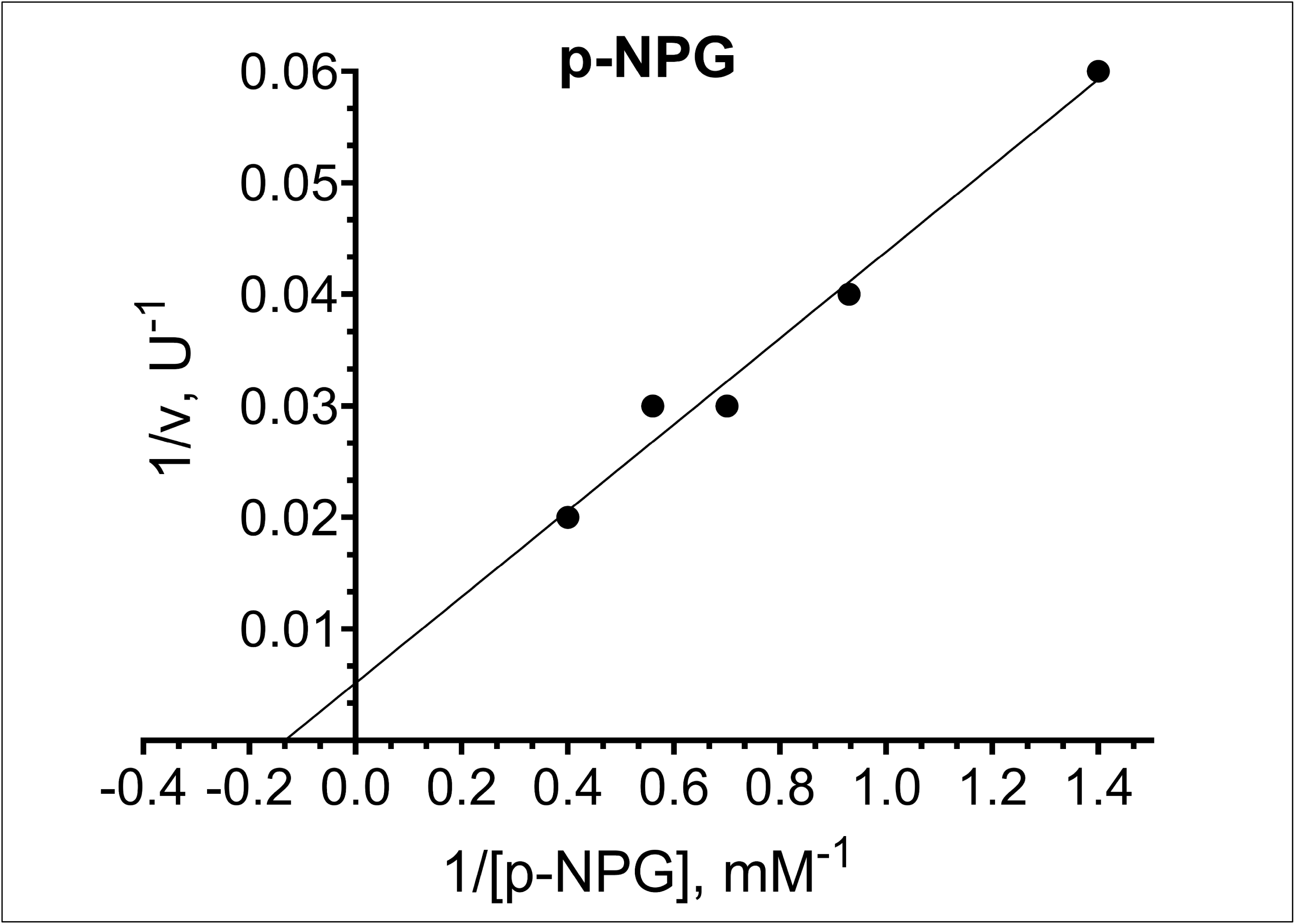
(**A**) The plot of relative activity (%) of Human Leukocyte Glucosylceramidase (GCase) in different concentrations of δ-gluconolactone **(B)** Lineweaver–Burk plot analysis of GCase inhibited by increasing the concentration of δ-gluconolactone. GCase activity was measured with *p*-nitrophenyl-β-D-glucopyranoside (PNPG) as the substrate, in the absence (no-inhibitor) or presence of the following concentration of δ-gluconolactone concentrations: (Inhibitor 1) 0.0128 mM; (Inhibitor 2) 0.0642 mM. The intercept of the plots indicates competitive inhibition for δ-gluconolactone (See Appendix A2 for inlet of intersection).

**Table 2.** Kinetic parameters of Glucosylceramidase in different concentration of gluconolactone and glucose

| Gluconolactone | $I_0$ | $I_1$ | $I_2$ | Conclusion |
| --- | --- | --- | --- | --- |
| **Vmax and $V_{max}^{app}$** | 333.3 | 333.3 | 333.3 | $V_{max}$ unchanged |
| **Km and $K_m^{app}$** | 3.6 | 5.6 | 8.6 | The apparent $K_m$ increased |
| **$K_i$** |  | 0.023 | 0.119 | Competitive Inhibition |
| Glucose | $I_0$ | $I_1$ | $I_2$ | Conclusion |
| **Vmax and $V_{max}^{app}$** | 333.3 | 90.9 | 13.5 | $V_{max}$ decreased |
| **Km and $K_m^{app}$** | 8.9 | 2.6 | 0.27 | The apparent $K_m$ decreased |
| **$K_i$** |  | 2.09 | 1.94 | Uncompetitive Inhibition |

**Figure 3.**
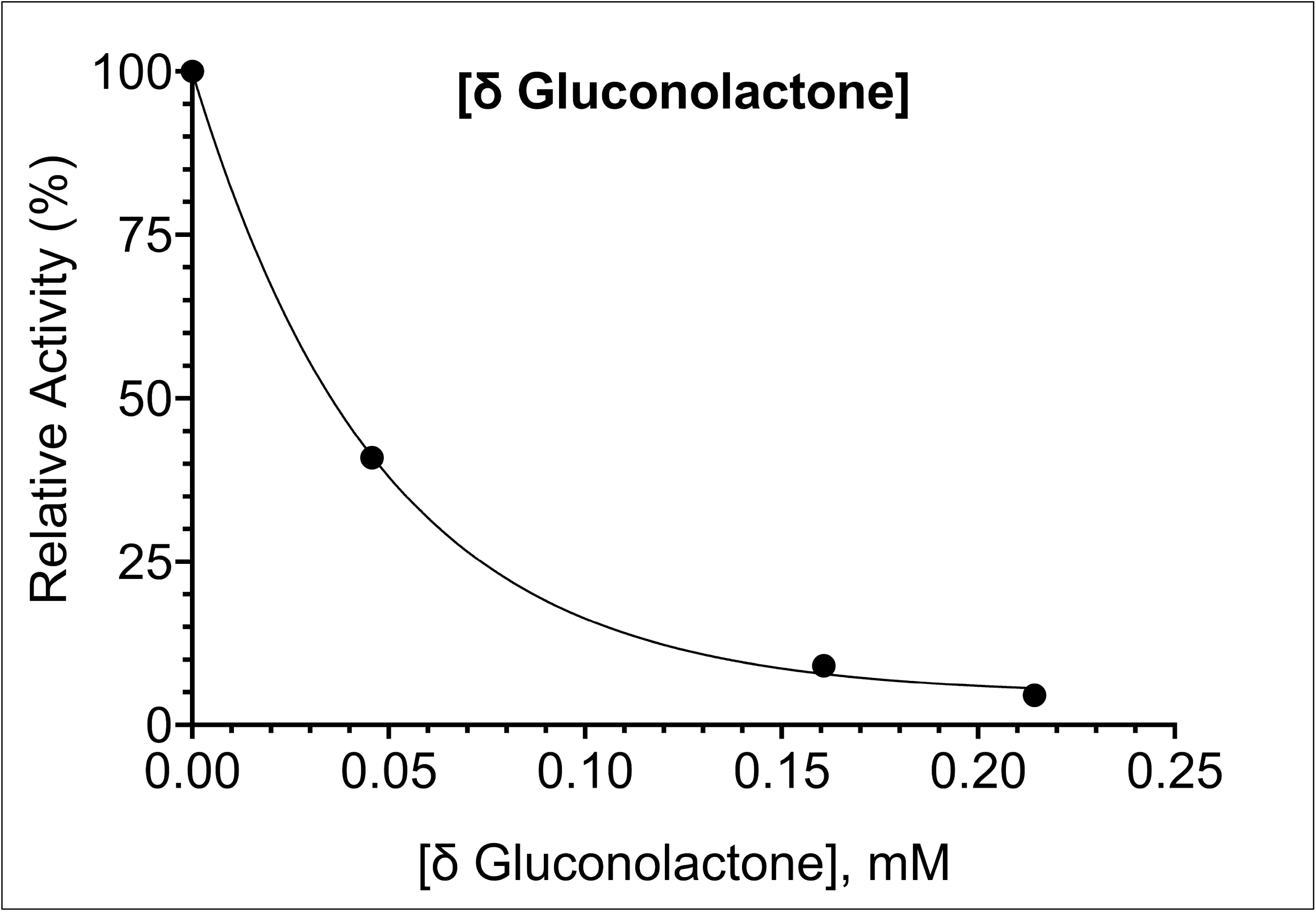
(**A**) The relative activity (%) curve of the Glucosylceramidase (GCase) in the presence of different glucose concentrations **(B)** Lineweaver–Burk plot analysis of the GCase inhibited by increasing the concentration of glucose. GCase activity was measured with *p*-nitrophenyl-β-D-glucopyranoside (PNPG) as the substrate, in the absence (no-inhibitor) or presence of the following concentration of glucose concentrations: (Inhibitor-1) 5.07 mM; (Inhibitor-2) 62.8 mM. Successful enzyme activity inhibition in an uncompetitive manner of glucose has been demonstrated. (See Appendix A2 for inlet of intersection).

## Abbreviations

GCase: Glucosylceramidase
GC: Glucosylceramide
GD: Gaucher disease
PD: Parkinson Disease
DLB: Dementia with Lewy bodies
*GBA1*: Glucosylceramidase beta acid 1
PNPG: *para*-Nitrophenyl-β-D-glucopyranoside
4-MUG: 4-methylumbelliferyl-β-D-glucopyranoside
IC_50_: Inhibitor concentration at which the rate is decreased by half

---

**Figure A1.**
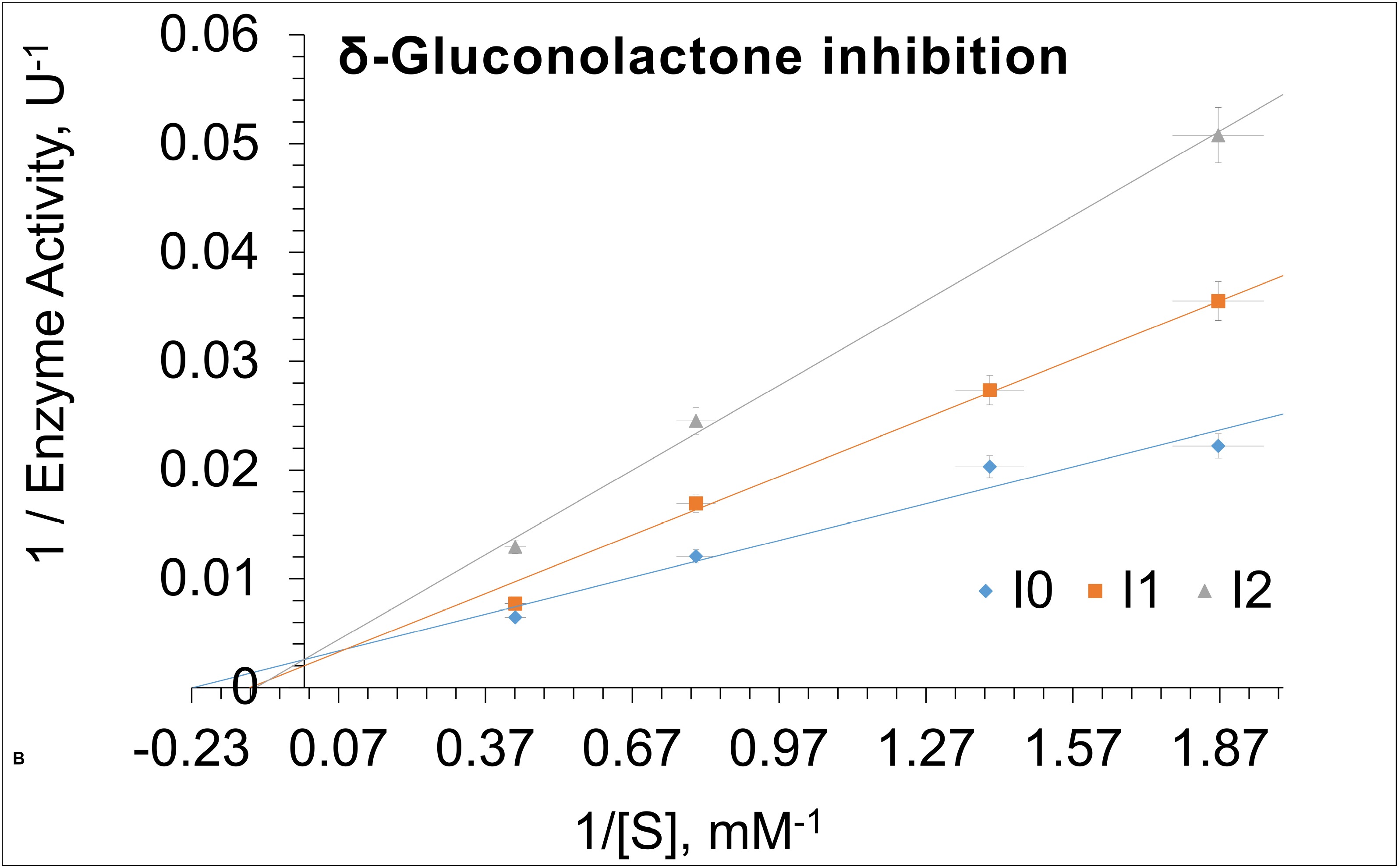
Glucosylceramidase (GCase) enzyme stability in leukocyte homogenate storage at −20°C. Enzyme activity at day 0 was taken as 100%, and the remaining results were normalized according to this value.

**Figure A2.**
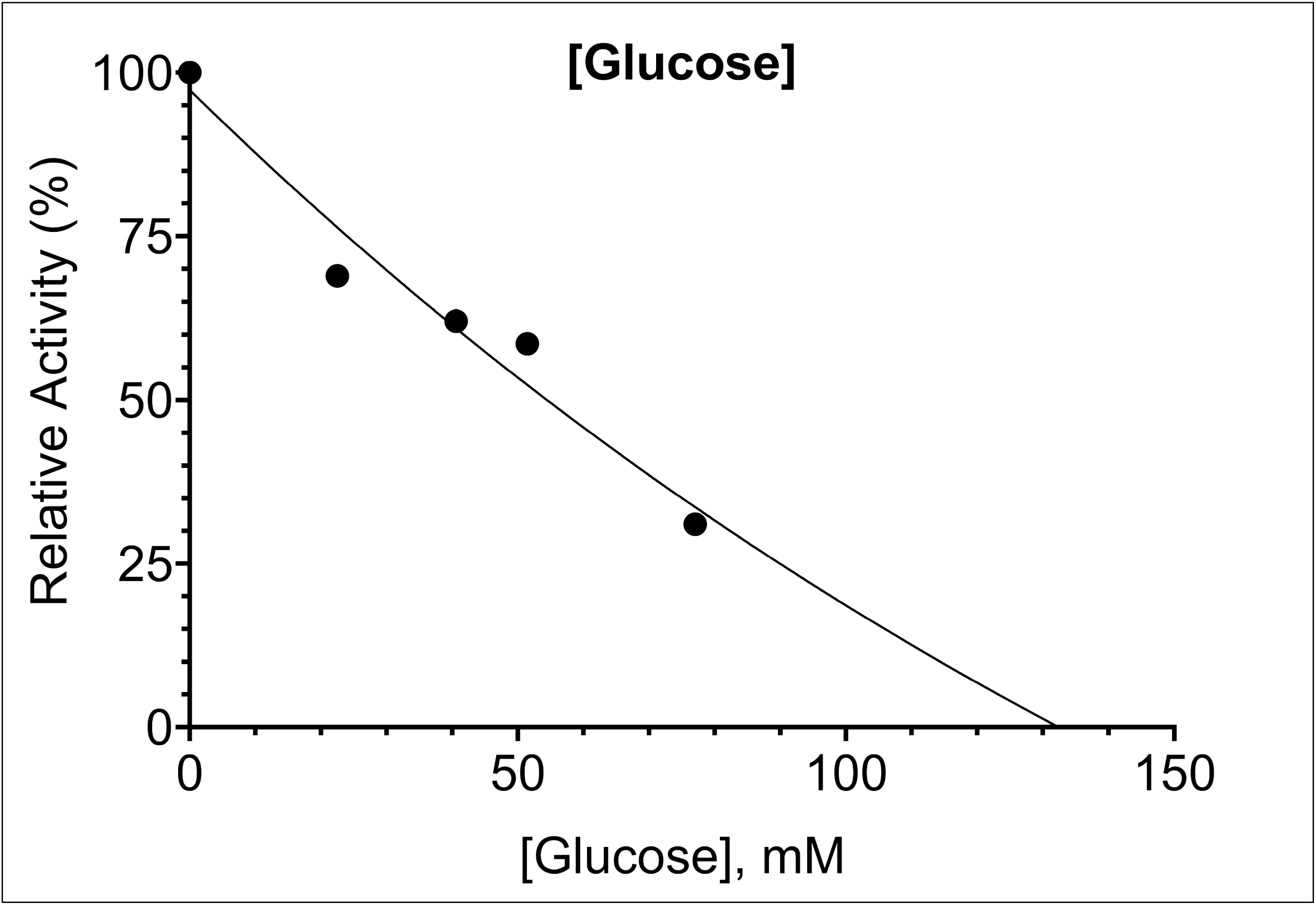
Inlet of intersection for Lineweaver–Burk plot analysis in Figures 2B and 3B for the GCase inhibition by increasing concentration of (**A**) δ-gluconolactone (**B**) glucose.

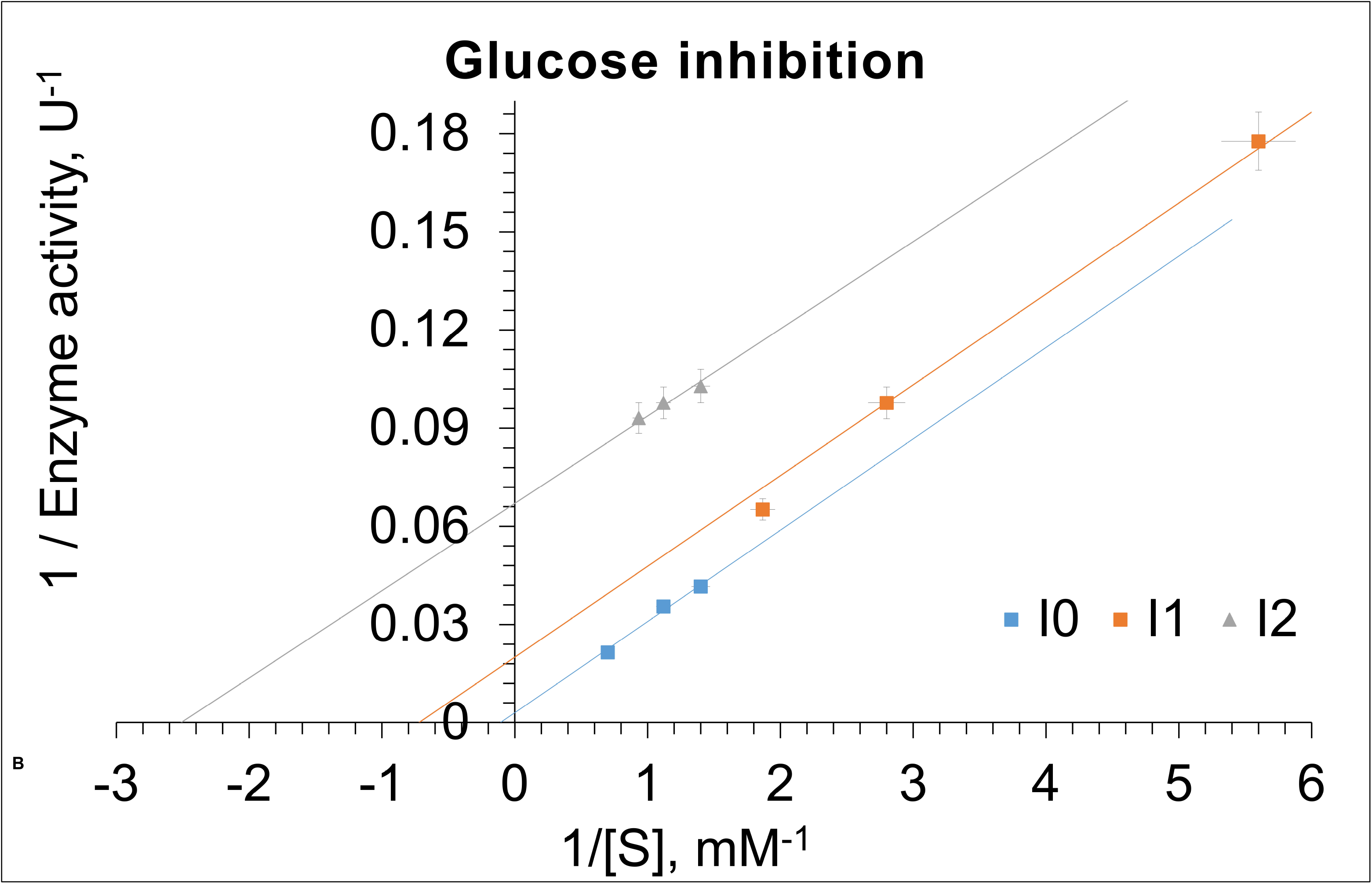

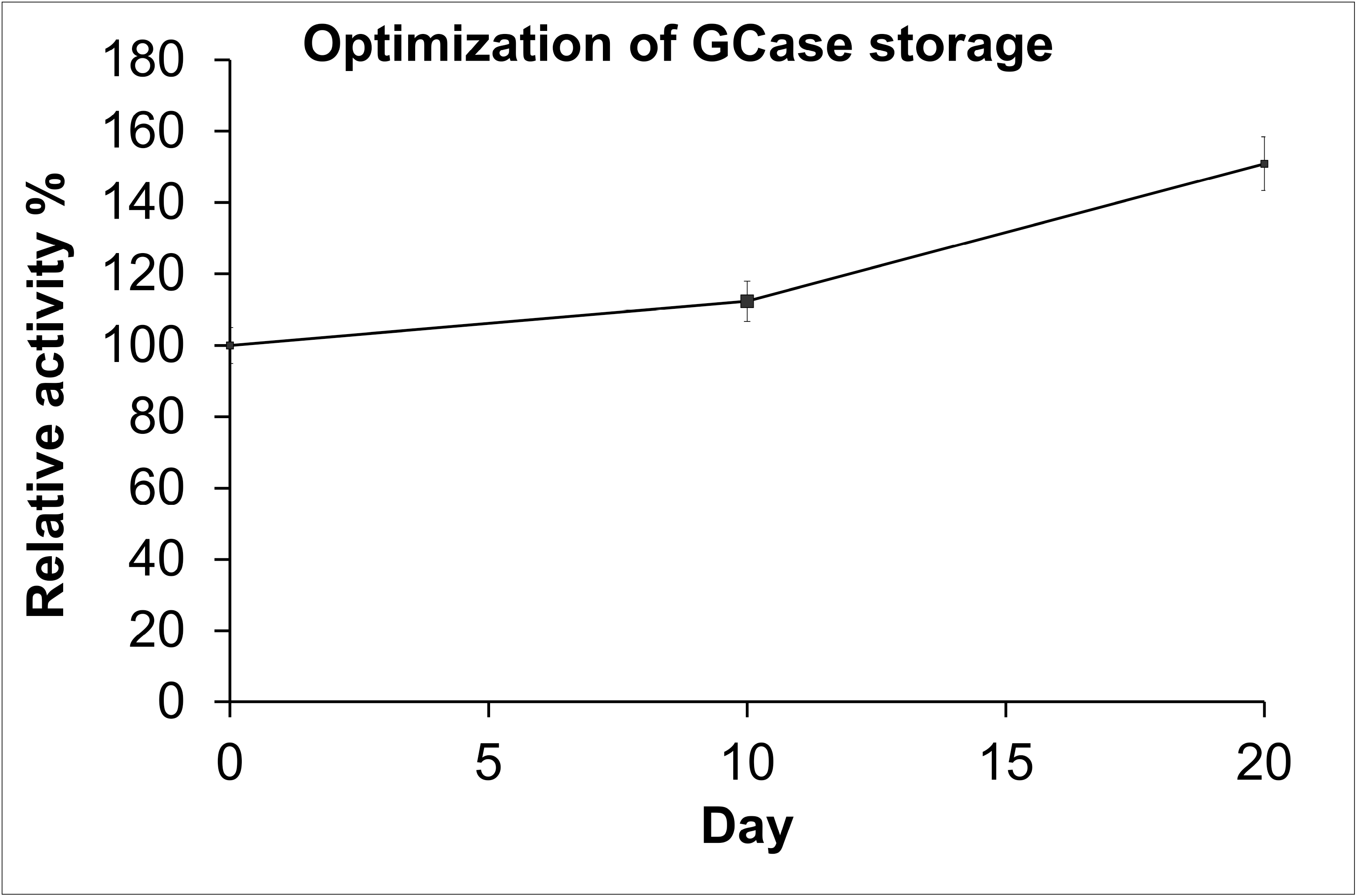

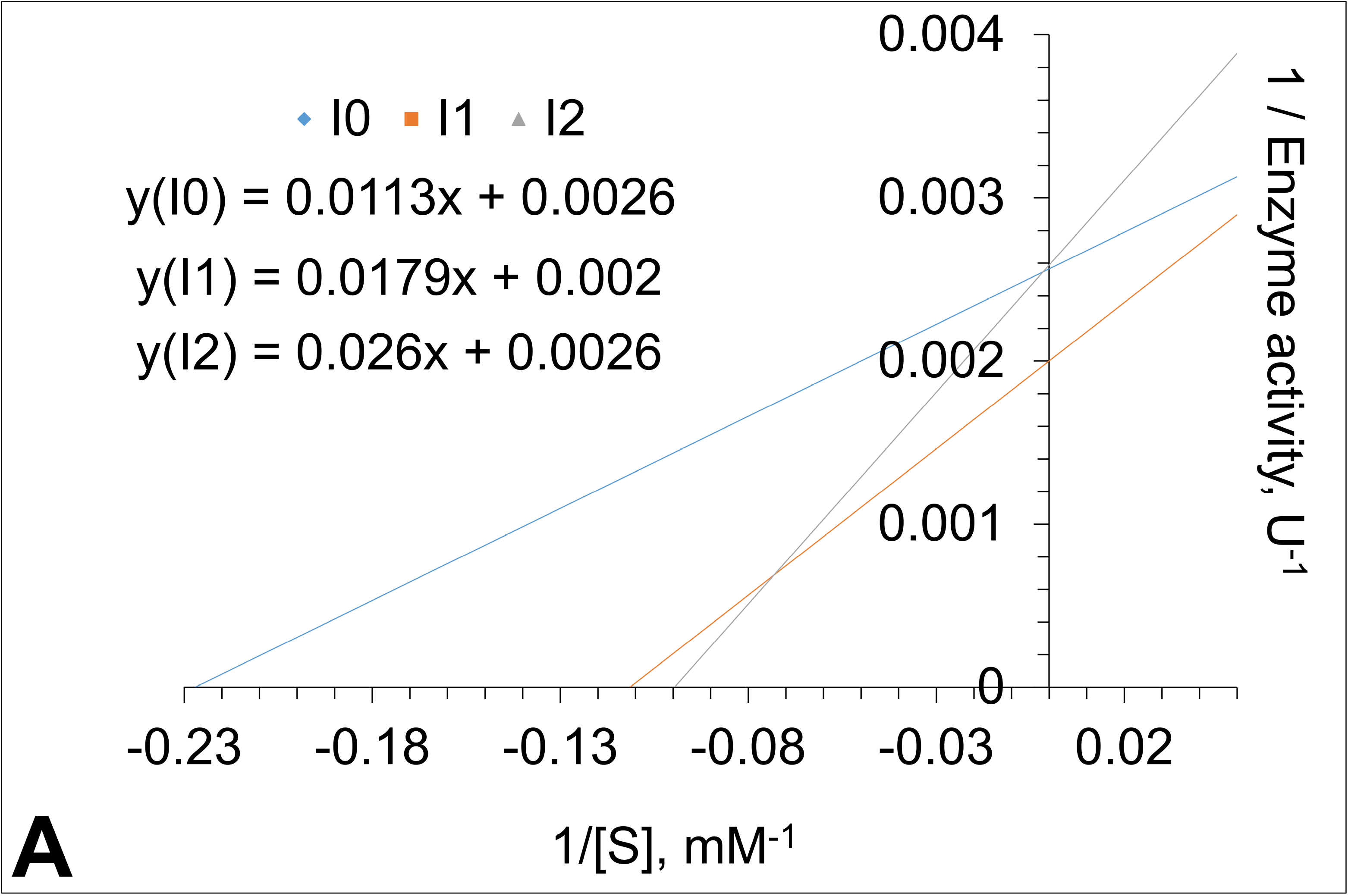

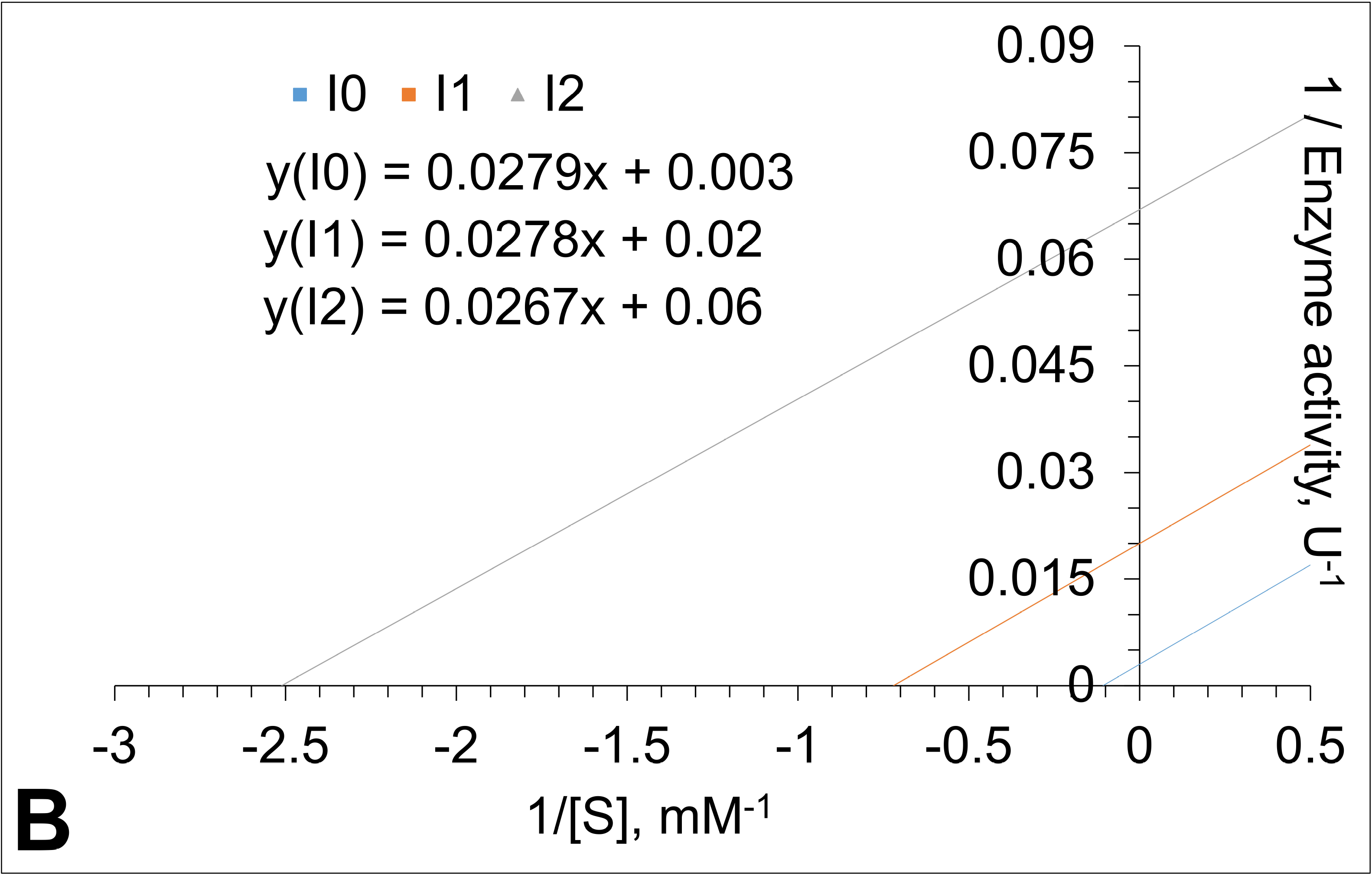

## REFERENCES

1. Smith L, Mullin S, Schapira AHV (2017) Insights into the structural biology of Gaucher disease. Exp Neurol 298:180–190. https://doi.org/10.1016/j.expneurol.2017.09.010

2. Beutler E (1992) Gaucher disease: new molecular approaches to diagnosis and treatment. Science 256:794–799. https://doi.org/10.1126/science.256.5058.794

3. Magalhaes J, Gegg ME, Migdalska-Richards A, et al (2016) Autophagic lysosome reformation dysfunction in glucocerebrosidase deficient cells: relevance to Parkinson disease. Hum Mol Genet 25:3432–3445. https://doi.org/10.1093/hmg/ddw185

4. Yap TL, Jiang Z, Heinrich F, et al (2015) Structural Features of Membrane-bound Glucocerebrosidase and α-Synuclein Probed by Neutron Reflectometry and Fluorescence Spectroscopy. J Biol Chem 290:744–754. https://doi.org/10.1074/jbc.M114.610584

5. Alroy J, Lyons JA (2014) Lysosomal storage diseases. J Inborn Errors Metab Screen 2:2326409813517663

6. Grabowski GA, Gaft S, Horowitz M, Kolodny EH (1990) Acid β-Glucosidase: Enzymology and Molecular Biology of Gaucher Diseas. Crit Rev Biochem Mol Biol 25:385–414

7. Glew RH, Basu A, LaMarco KL, Prence EM (1989) Mammalian glucocerebrosidase: implications for Gaucher’s disease. In: Pathology Reviews· 1989. Springer, pp 3–23

8. Kim H-J, Jeon B, Song J, et al (2016) Leukocyte glucocerebrosidase and β-hexosaminidase activity in sporadic and genetic Parkinson disease. Parkinsonism Relat Disord 23:99–101. https://doi.org/10.1016/j.parkreldis.2015.12.002

9. Goker-Alpan O, Giasson BI, Eblan MJ, et al (2006) Glucocerebrosidase mutations are an important risk factor for Lewy body disorders. Neurology 67:908–910. https://doi.org/10.1212/01.wnl.0000230215.41296.18

10. Karatas M, Turan Y, Kurtovic-Kozaric A, Dogan S (2016) Analysis of Gaucher Disease Responsible Genes in Colorectal Adenocarcinoma. J Biom Biostat 7:. https://doi.org/10.4172/2155-6180.1000314

11. Cormand B, Montfort M, Chabás A, et al (1997) Genetic fine localization of the β-glucocerebrosidase (GBA) and prosaposin (PSAP) genes: implications for Gaucher disease. Hum Genet 100:75–79. https://doi.org/10.1007/s004390050468

12. Sheth J, Bhavsar R, Mistri M, et al (2019) Gaucher disease: single gene molecular characterization of one-hundred Indian patients reveals novel variants and the most prevalent mutation. BMC Med Genet 20:31. https://doi.org/10.1186/s12881-019-0759-1

13. Dogan S, Kurtovic-Kozaric A, Karlı, Gunay (2016) The Detection of Extremely High and Low Expressed Genes by EGEFAlgorithm in Invasive Breast Cancer. https://doi.org/10.4172/2155-6180.1000286

14. Meivar-Levy I, Horowitz M, Futerman AH (1994) Analysis of glucocerebrosidase activity using N-(1-[14C]hexanoyl)-D-erythroglucosylsphingosine demonstrates a correlation between levels of residual enzyme activity and the type of Gaucher disease. Biochem J 303:377–382

15. Grace ME, Newman KM, Scheinker V, et al (1994) Analysis of human acid beta-glucosidase by site-directed mutagenesis and heterologous expression. J Biol Chem 269:2283–2291

16. Glew RH, Basu A, LaMarco KL, Prence EM (1989) Mammalian glucocerebrosidase: implications for Gaucher’s disease. In: Pathology Reviews· 1989. Springer, pp 3–23

17. Spahiu E (2015) ATR-FTIR Evaluation of Structural and Functional Changes on Murine Macrophage Cells Upon Activation and Suppression by Immuno-Therapeutic Oligodeoxynucleotides. Thesis, Middle East Technical University

18. Beutler E, Demina A, Gelbart T (1994) Glucocerebrosidase mutations in Gaucher disease. Mol Med 1:82

19. A Z, A K, T G, et al (1992) Gaucher disease. Clinical, laboratory, radiologic, and genetic features of 53 patients. Medicine (Baltimore) 71:337–353

20. Dogan S, Mermer E (2017) Comparison of the Hemoglobin Amount between Old and Young Persons in Bosnia and Herzegovina. J Biom Biostat 08: https://doi.org/10.4172/2155-6180.1000337

21. Pastores GM, Hughes DA (1993) Gaucher Disease. In: Adam MP, Ardinger HH, Pagon RA, et al (eds) GeneReviews®. University of Washington, Seattle, Seattle (WA)

22. Naphatsamon U, Ohashi T, Misaki R, Fujiyama K (2018) The Production of Human β-Glucocerebrosidase in Nicotiana benthamiana Root Culture. Int J Mol Sci 19:1972. https://doi.org/10.3390/ijms19071972

23. Bešić L, Ašić A, Muhović I, et al (2017) Purification and Characterization of β-Glucosidase from Brassica oleracea. J Food Process Preserv 41:

24. De Fost M, Aerts JM, Hollak CE (2003) Gaucher disease: from fundamental research to effective therapeutic interventions. Neth J Med 61:3–8

25. Pastores GM, Hughes DA (1993) Gaucher Disease. In: Adam MP, Ardinger HH, Pagon RA, et al (eds) GeneReviews®. University of Washington, Seattle, Seattle (WA)

26. Yang N-Y, Lee Y-N, Lee H-J, et al (2013) Glucocerebrosidase, a new player changing the old rules in Lewy body diseases. Biol Chem 394:807–818. https://doi.org/10.1515/hsz-2012-0322

27. Raghavan SS, Topol J, Kolodny EH (1980) Leukocyte β-glucosidase in homozygotes and heterozygotes for Gaucher disease. Am J Hum Genet 32:158

28. Huang C, Freter C (2015) Lipid metabolism, apoptosis and cancer therapy. Int J Mol Sci 16:924–949

29. Glew RH, Basu A, Prence E, et al (1986) The Effects of Acidic Lipids and Heat-Stable Factor on the Physical-Chemical and Kinetic Properties of Glucocerebrosidase. In: Enzymes of Lipid Metabolism II. Springer, pp 361–370

30. Magalhaes J, SáMiranda MC, Pinto R, et al (1984) Sodium taurocholate effect on β-glucosidase activity: a new approach for identification of Gaucher disease using the synthetic substrate and leucocytes. Clin Chim Acta 141:111–118

31. Wattiaux R, De Duve C (1956) Tissue fractionation studies. 7. Release of bound hydrolases by means of Triton X-100. Biochem J 63:606

32. Glew RH, Rosenthal MD (2006) Clinical Studies in Medical Biochemistry. Oxford University Press

33. Butcher BA, Gopalan V, Lee RE, et al (1989) Use of 4-heptylumbelliferyl-β-d-glucoside to identify Gaucher’s disease heterozygotes. Clin Chim Acta 184:235–242

34. Daniels LB, Glew RH (1982) beta-Glucosidase assays in the diagnosis of Gaucher’s disease. Clin Chem 28:569–577

35. Ho MW (1973) Identity of ‘acid’β-glucosidase and glucocerebrosidase in human spleen. Biochem J 136:721–729

36. Ho MW, O’Brien JS (1971) Gaucher’s Disease: Deficiency of “cid” β-Glucosidase and Reconstitution of Enzyme Activity In Vitro. Proc Natl Acad Sci 68:2810–2813

37. Hyun JC, Misra RS, Greenblatt D, Radin NS (1975) Synthetic inhibitors of glucocerebroside β-glucosidase. Arch Biochem Biophys 166:382–389

38. Lieberman RL (2011) A guided tour of the structural biology of Gaucher disease: Acid-β-glucosidase and saposin C. Enzyme Res 2011:

39. Michelin K, Wajner A, Bock H, et al (2005) Biochemical properties of β-glucosidase in leukocytes from patients and obligated heterozygotes for Gaucher disease carriers. Clin Chim Acta 362:101–109

40. Michelin K, Wajner A, Goulart L da S, et al (2004) Biochemical study on β-glucosidase in individuals with Gaucher’s disease and normal subjects. Clin Chim Acta 343:145–153

41. Peters SP, Coyle P, Glew RH (1976) Differentiation of β-glucocerebrosidase from β-glucosidase in human tissues using sodium taurocholate. Arch Biochem Biophys 175:569–582

42. Peters SP, Lee RE, Glew RH (1975) A microassay for Gaucher’s disease. Clin Chim Acta 60:391–396

43. Strasberg PM, Lowden JA (1982) The assay of glucocerebrosidase activity using the natural substrate. Clin Chim Acta 118:9–20

44. Wenger DA, Clark C, Sattler M, Wharton C (1978) Synthetic substrate \s s-glucosidase activity in leukocytes: A reproducible method for the identification of patients and carriers of Gaucher’s disease. Clin Genet 13:145–153

45. Yu-Bin C, Glew RH, Diven WF, Lee RE (1980) Comparison of various β-glucosidase assays used to diagnose Gaucher’s disease. Clin Chim Acta 105:41–50

46. Fishman WH, Springer B, Brunetti R (1948) Application of an improved glucuronidase assay method to the study of human blood β-glucuronidase. J Biol Chem 173:449–456

47. Kato K, Yoshida K, Tsukamoto H, et al (1960) Synthesisof p-Nitrophenyl β-D-Glucopyranosiduronic Acid and Its Utilization as a Substrate for the Assay of β-Glucuronidase Activity. Chem Pharm Bull (Tokyo) 8:239–242

48. Szasz G (1967) Comparison between p-nitrophenyl glucuronide and phenolphthalein glucuronide as substrates in the assay of β-glucuronidase. Clin Chem 13:752–759

49. Beutler E, Kuhl W (1970) The diagnosis of the adult type of Gaucher’s disease and its carrier state by demonstration of deficiency of β-glucosidase activity in peripheral blood leukocytes. J Lab Clin Med 76:747–755

50. Grabowski GA, Dagan A (1984) Human lysosomal β-glucosidase: Purification by affinity chromatography. Anal Biochem 141:267–279

51. Ho MW, Seck J, Schmidt D, et al (1972) Adult Gaucher’s disease: kindred studies and demonstration of a deficiency of acid beta-glucosidase in cultured fibroblasts. Am J Hum Genet 24:37

52. Maret A, Salvayre R, Potier M, et al (1988) Comparison of Human Membrane-Bound β-Glucosidases: Lysosomal Glucosylceramide-β-Glucosidase and Non-Specific β-Glucosidase. In: Lipid Storage Disorders. Springer, pp 57–61

53. Paal K, Ito M, Withers SG (2004) Paenibacillus sp. TS12 glucosylceramidase: kinetic studies of a novel sub-family of family 3 glycosidases and identification of the catalytic residues. Biochem J 378:141–149. https://doi.org/10.1042/BJ20031028

54. Karatas M, Dogan S, Spahiu E, A Ašić, L Bešić, Y Turan (2020) Preparation of leukocytes by differential lysis of erythrocytes. Protocols.io https://doi.org/10.17504/protocols.io.bcv2iw8e

55. Kara HE, Sinan S, Turan Y (2011) Purification of beta-glucosidase from olive (Olea europaea L.) fruit tissue with specifically designed hydrophobic interaction chromatography and characterization of the purified enzyme. J Chromatogr B 879:1507–1512. https://doi.org/10.1016/j.jchromb.2011.03.036

56. Bradford MM (1976) A rapid and sensitive method for the quantitation of microgram quantities of protein utilizing the principle of protein-dye binding. Anal Biochem 72:248–254

57. Raynal B, Lenormand P, Baron B, et al (2014) Quality assessment and optimization of purified protein samples: why and how? Microb Cell Factories 13:. https://doi.org/10.1186/s12934-014-0180-6

58. Raynal B, Lenormand P, Baron B, et al (2014) Quality assessment and optimization of purified protein samples: why and how? Microb Cell Factories 13:. https://doi.org/10.1186/s12934-014-0180-6

59. Xia W, Xu X, Qian L, et al (2016) Engineering a highly active thermophilic β-glucosidase to enhance its pH stability and saccharification performance. Biotechnol Biofuels 9:147. https://doi.org/10.1186/s13068-016-0560-8

60. Tan YL, Genereux JC, Pankow S, et al (2014) ERdj3 Is an Endoplasmic Reticulum Degradation Factor for Mutant Glucocerebrosidase Variants Linked to Gaucher’s Disease. Chem Biol 21:967–976. https://doi.org/10.1016/j.chembiol.2014.06.008

61. Grabowski GA, Gatt S, Kruse J, Desnick RJ (1984) Human lysosomal β-glucosidase: kinetic characterization of the catalytic, aglycon, and hydrophobic binding sites. Arch Biochem Biophys 231:144–157

62. Ašić A, Bešić L, Muhović I, et al (2015) Purification and characterization of β-glucosidase from Agaricus bisporus (white button mushroom). Protein J 34:453–461

63. Bešić L, Ašić A, Muhović I, et al (2017) Purification and Characterization of β-Glucosidase from Brassica oleracea. J Food Process Preserv 41:

64. Teugjas H, Väljamäe P (2013) Selecting β-glucosidases to support cellulases in cellulose saccharification. Biotechnol Biofuels 6:105

65. Uchiyama T, Miyazaki K, Yaoi K (2013) Characterization of a novel β-glucosidase from a compost microbial metagenome with strong transglycosylation activity. J Biol Chem 288:18325–18334

66. Turan Y, Zheng M (2005) Purification and characterization of an intracellular β-glucosidase from the methylotrophic yeast Pichia pastoris. Biochem Mosc 70:1363–1368

67. Chan CS, Sin LL, Chan K-G, et al (2016) Characterization of a glucose-tolerant β-glucosidase from Anoxybacillus sp. DT3-1. Biotechnol Biofuels 9:174

68. Pei J, Pang Q, Zhao L, et al (2012) Thermoanaerobacterium thermosaccharolyticum β-glucosidase: a glucose-tolerant enzyme with high specific activity for cellobiose. Biotechnol Biofuels 5:31

